# Barcode aware adaptive sampling for GridION and PromethION Oxford Nanopore sequencers

**DOI:** 10.1101/2021.12.01.470722

**Authors:** Alexander Payne, Rory Munro, Nadine Holmes, Christopher Moore, Matt Carlile, Matthew Loose

## Abstract

Adaptive sampling enables selection of individual DNA molecules from sequencing libraries, a unique property of nanopore sequencing. Here we develop our adaptive sampling tool readfish to become “barcode-aware” enabling selection of different targets within barcoded samples or filtering out individual barcodes. We show that multiple human genomes can be assessed for copy number and structural variation on a single sequencing flow cell using sample specific customised target panels on both GridION and PromethION devices.

## Main Text

Adaptive sampling is the process by which individual DNA molecules within a library can be dynamically selected for sequencing, a property unique to Oxford Nanopore Technologies (ONT) sequencers ^1^. Recently we developed readfish, which uses real-time base calling to analyse read data as molecules are being sequenced ^2^. Using readfish, it is possible to enrich target regions of human genomes as well as manipulate sequencing coverage of metagenomic samples ^2–4^. Here we show this method can be extended to enable the use of barcoded samples with readfish. This allows for individual barcodes to be switched off during a run or enables the use of targets specific to each barcode. We also demonstrate that this method can be used with Oxford Nanopore PromethION devices to enable high coverage over multiple targets from different genomes in a single experiment.

An advantage of sequence based approaches to adaptive sampling is that existing tools, such as barcode demultiplexers, can be easily incorporated into the readfish workflow. Although signal based methods to identify barcodes exist, no sufficiently fast methods are currently available ^5^. We therefore adapted our existing readfish pipeline to be compatible with built-in Guppy demultiplexing (ONT) and incorporated barcode classifications into the data readfish uses to make a decision about sequencing or rejecting a read. To test adaptive sampling on the PromethION device we additionally used Guppy’s ability to generate read mappings.

An important consideration in adaptive sampling is the duration of signal data needed for accurate mapping of a read fragment. Previously we used chunks of 0.4 seconds of data ^2^, (roughly 1,600 samples) but reasoned the inclusion of additional barcode sequence at the start of each read would require additional data. To test this we took a set of reads (see methods) and sampled signals from the start of each representing the data seen when running adaptive sampling. We then used a variety of base caller models (see methods) and two signal alignment tools, Uncalled and Sigmap, to analyse mappings from each of these synthetic reads and methods ^3,6^. As readfish uses the start coordinate of a mapping to determine if a read is on target, we compared the predicted mapping coordinate with that from the full length read (high accuracy mode - HAC). A correct mapping is defined as one where the start mapping coordinates are within 100 bases of one another. We found that 3,600 samples (or 0.8 seconds of data) was sufficient to correctly place reads (F1>0.9, fast model) (Figure 1A). Similarly, this same window also enabled appropriate barcode mapping accuracy (F1>0.9, fast model) (Figure 1B). Therefore we configured our GridION experiments to use data in chunks of 0.8 seconds (3,200 samples of data). For PromethION, we remained with the default value of 1 second of data so as to reduce the total analysis load due to the higher capacity flow cell. This is an area for future optimisation.

**Figure 1.**
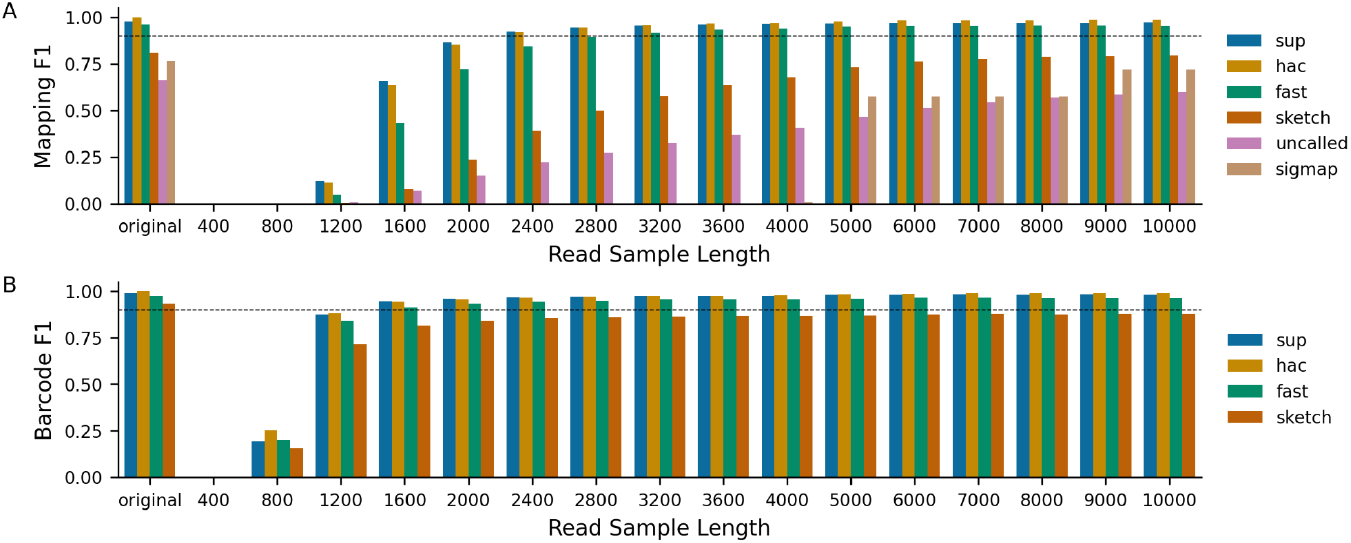
Comparison of base callers, alignments and barcode classifications. A set of pass reads derived from chromosome 15 and 17 targets from the truSight Fusion panel were generated. A) Reads were base called using the super accuracy (sup), high accuracy (hac), fast (fast) or sketch (sketch) models of guppy and mapped to a synthetic genome containing only the target regions for those read targets. These same reads were also mapped using the signal aligners Uncalled and Sigmap. Truth was defined as the start mapping coordinate for the full length read (original). Read fragments were scored as mapping correctly if the start mapping coordinates were within 50 bases of the true start mapping position. 0.9 F1 is exceeded at 0.8 seconds of data (3200 samples) for the fast model. B) F1 score as measured by concordance in barcode identified where truth is the HAC model.

Readfish can be configured to handle barcodes in two ways. For simple experiments, the user can identify a list of barcodes to be either rejected or accepted. In this way users can exclude or include a subset of barcodes on a sequencing run (Figure 2A). For more complex experiments, the user can configure a set of targets for each individual barcode in a library and so sequence specific regions from each. For example, a cancer gene panel for sample A, a developmental disorders panel for sample B and a neuropathology panel for sample C. Figure 2B illustrates a simple barcoded sample where different regions of a bacterial genome are selected on each barcode in real-time. As readfish maps using sequences it is only limited by available memory and easily handles gigabase genomes. In addition there is no requirement for each sample to be from the same organism and so readfish can target multiple references. To simplify creation and dynamic update of readfish configuration files, we provide a set of command line tools to configure options for multiple barcodes (https://github.com/looselab/readfish-tools).

**Figure 2.**
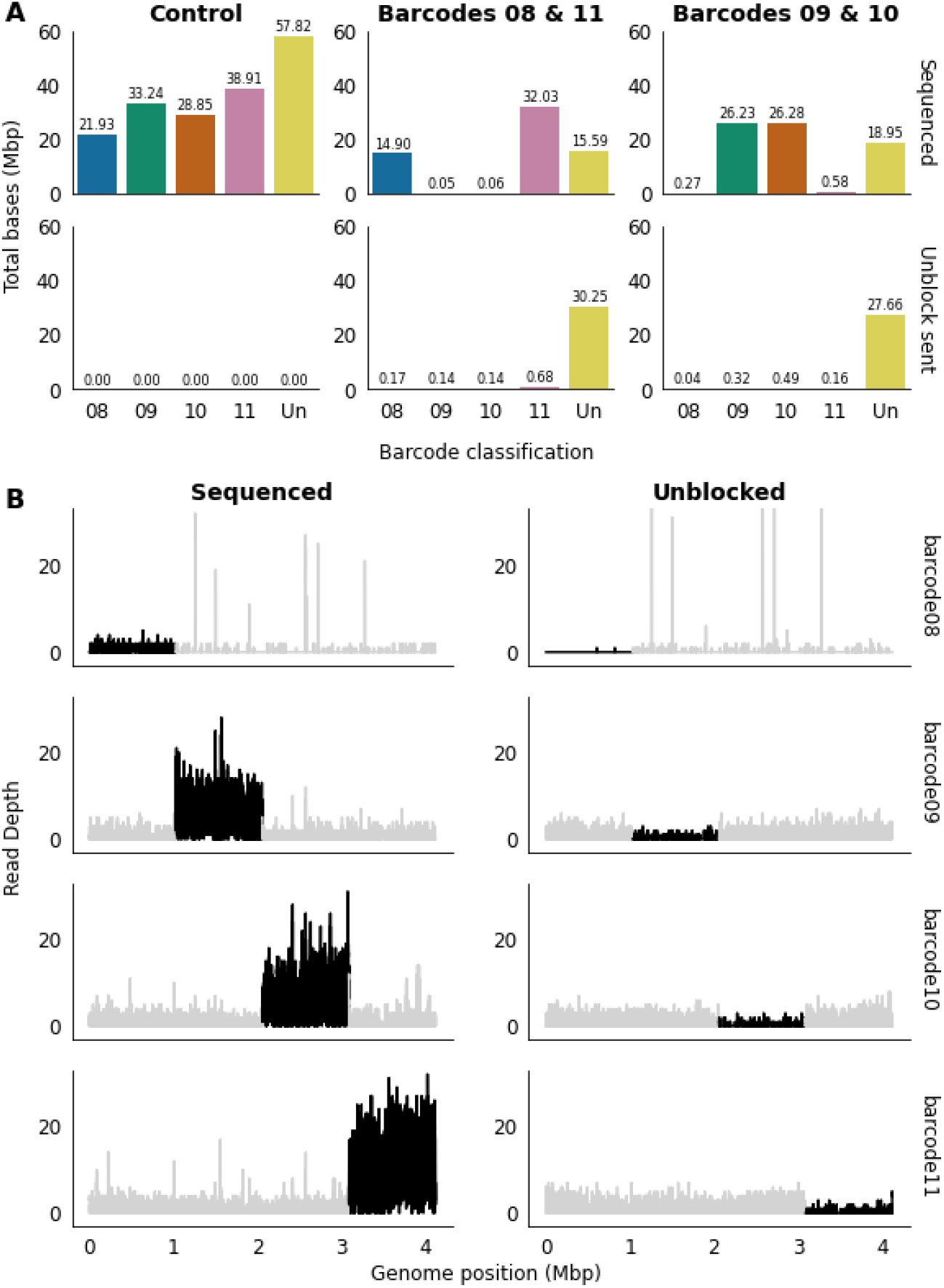
Naive barcoding selective sequencing. A) Demonstration of “switching off” individual barcodes from a sequencing library. Selected barcodes identified in the panel titles. Top row shows sequenced reads, lower panel shows the rejected or unblocked reads. As barcoding both ends is used to specify barcode, all rejected reads become unclassified (Un) by default. B) Switching the mode of operation for readfish from simple barcode rejection to differential targets. Sample shown is *Clostridioides difficile*. Targeted regions are shown in black. Unfortunately barcode08 was under-represented. in this sample, leading to low sequenced read count.

To test the performance of this approach, we used three previously described cell lines: GM12878, from the Utah/CEPH pedigree; NB4, a cell line carrying a fusion between PML and RARA representing an acute promyelocytic leukaemia (APL); and 22Rv1, a prostate cancer derived cell line containing significant chromosomal abnormalities ^7–9^. For each sample, we chose a specific gene panel. GM12878 was targeted using a panel defined by the gene list in the commercially available TruSight 170 Tumor panel ^10^. As the NB4 cell line contains an APL fusion, we selected the TruSight RNA Fusion Panel ^11^. For the more complex 22Rv1 prostate cancer line we used the previously described COSMIC panel ^2,12^. Samples were barcoded and sequenced on a single flow cell, and run for 72 hours (see methods). Every 24 hours the flow cell was washed with nuclease flush and another aliquot of the library loaded ^2^. On GridION, in a single experiment using a flow cell with 1,330 pores, 18.1 Gb of data were generated, with a total of 15 Gb successfully demultiplexed into barcoded data (Table 1). For PromethION testing, a single experiment using a flow cell with 6,960 pores generated a total of 78.2 Gb of data, of which 61 Gb were demultiplexed into barcoded data (Table 2).

**Table 1.**
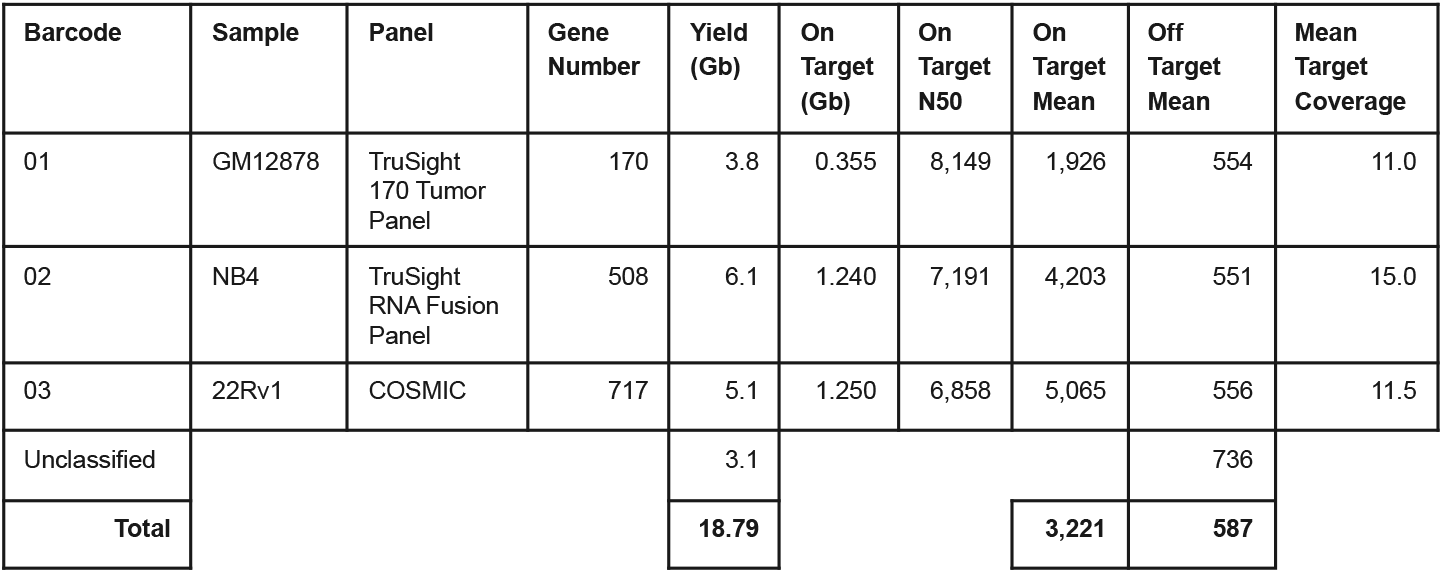
Sample Performance - GridION. Run metric performance per barcode and over the entire flow cell. Metrics are derived from real-time monitoring with minoTour.

**Table 2.**
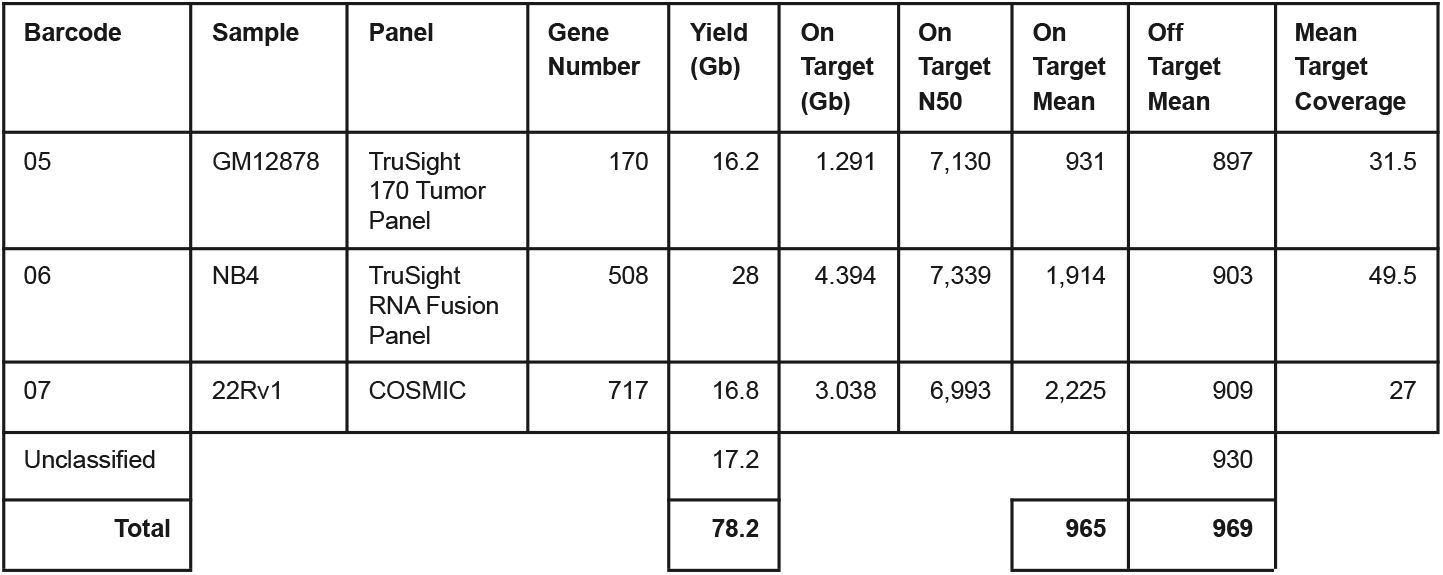
Sample Performance - PromethION. Run metric performance per barcode and over the entire flow cell. Metrics are derived from real-time monitoring with minoTour.

Across the whole GridION experiment, the on target read N50 was 7 kb, with the rejected read N50 being 579 bases, or approximately 1.3 seconds of sequencing. This results in mean read coverage on target regions of between 11x and 15x. Inspection of individual targets including BRCA1, NBR1, PML and RARA demonstrates the ability to specifically target unique regions on each sample (Figure 3, A-C). Current best practice for variant calling requires higher minimal depth than we achieve when looking at three samples. However, long range structural variants can be measured and so we used cuteSV ^13^ to analyse these three samples. As expected, multiple reads supporting the detection of a fusion between PML and RARA were detected in the NB4 cell line (Figure 4). In contrast, this rearrangement was not found in the 22Rv1 line. We cannot formally exclude the presence of this variant in GM12878 as neither PML or RARA were within the gene panel used for this cell line (Figures 3,4).

**Figure 3.**
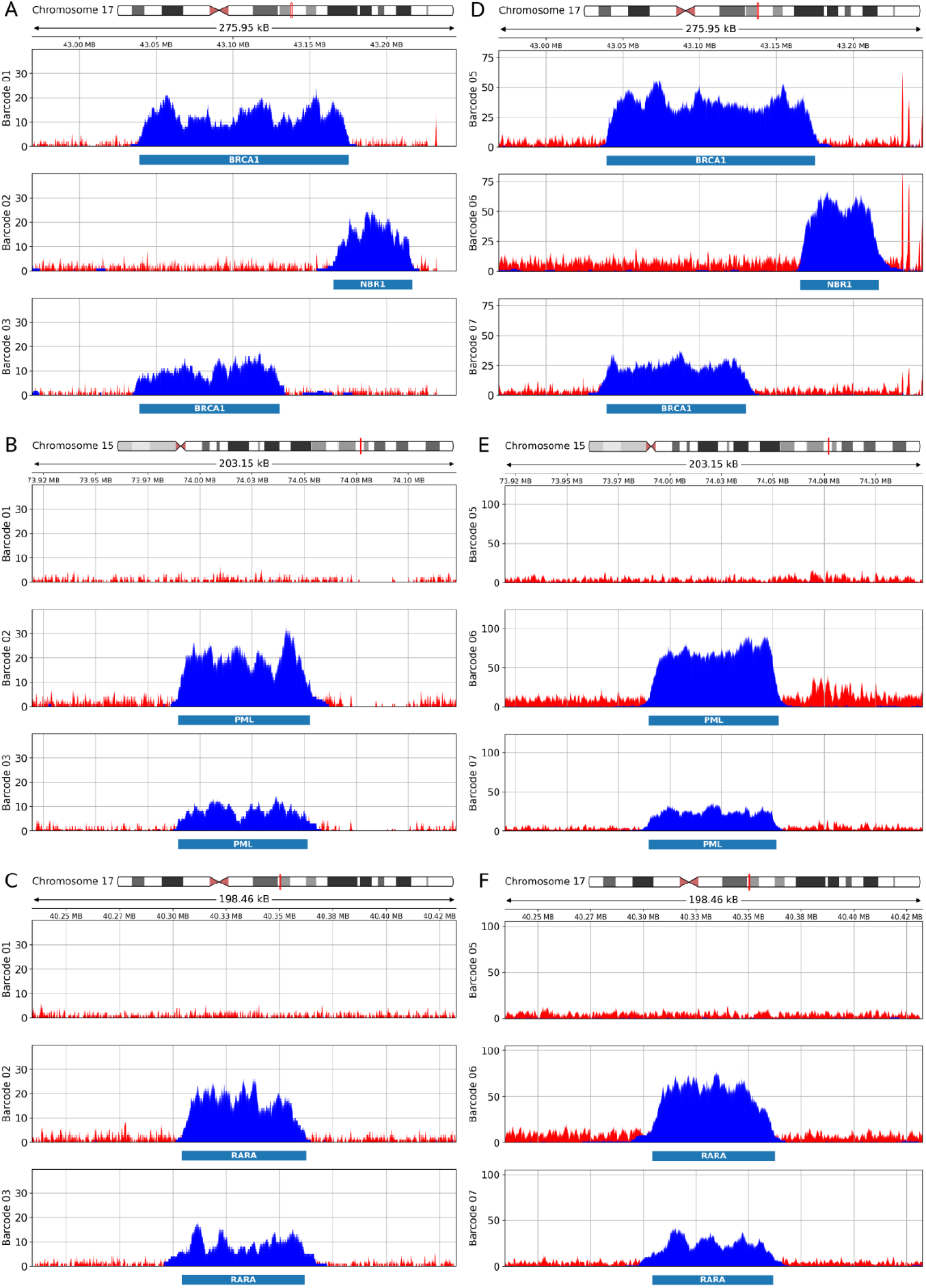
Target and barcode specific gene coverage. Illustration of coverage over each barcoded sample for each target in the panel. Blue is sequenced read coverage, red illustrates coverage of rejected reads. A-C) Data generated on a single GridION flowcell. A) shows coverage over BRCA1 and the adjacent gene NBR1. BRCA1 was a target for barcode 1 and 3, but not 2. The targeted regions are illustrated below the coverage plots. Note that the region representing BRCA1 differs in barcode 1 and 3 by design. NBR1 was only targeted on barcode 2. B) and C) illustrate coverage over PML and RARA respectively, which were only targeted on barcodes 2 and 3. D-F) Data generated on a single PromethION flowcell. Samples and targets as in A-C, but this library used barcodes 5,6 and 7. Barcode 01 and 05 are NA12878, Barcode 02 and 06 are NB4 and Barcode 03 and 07 are 22Rv1.

**Figure 4.**
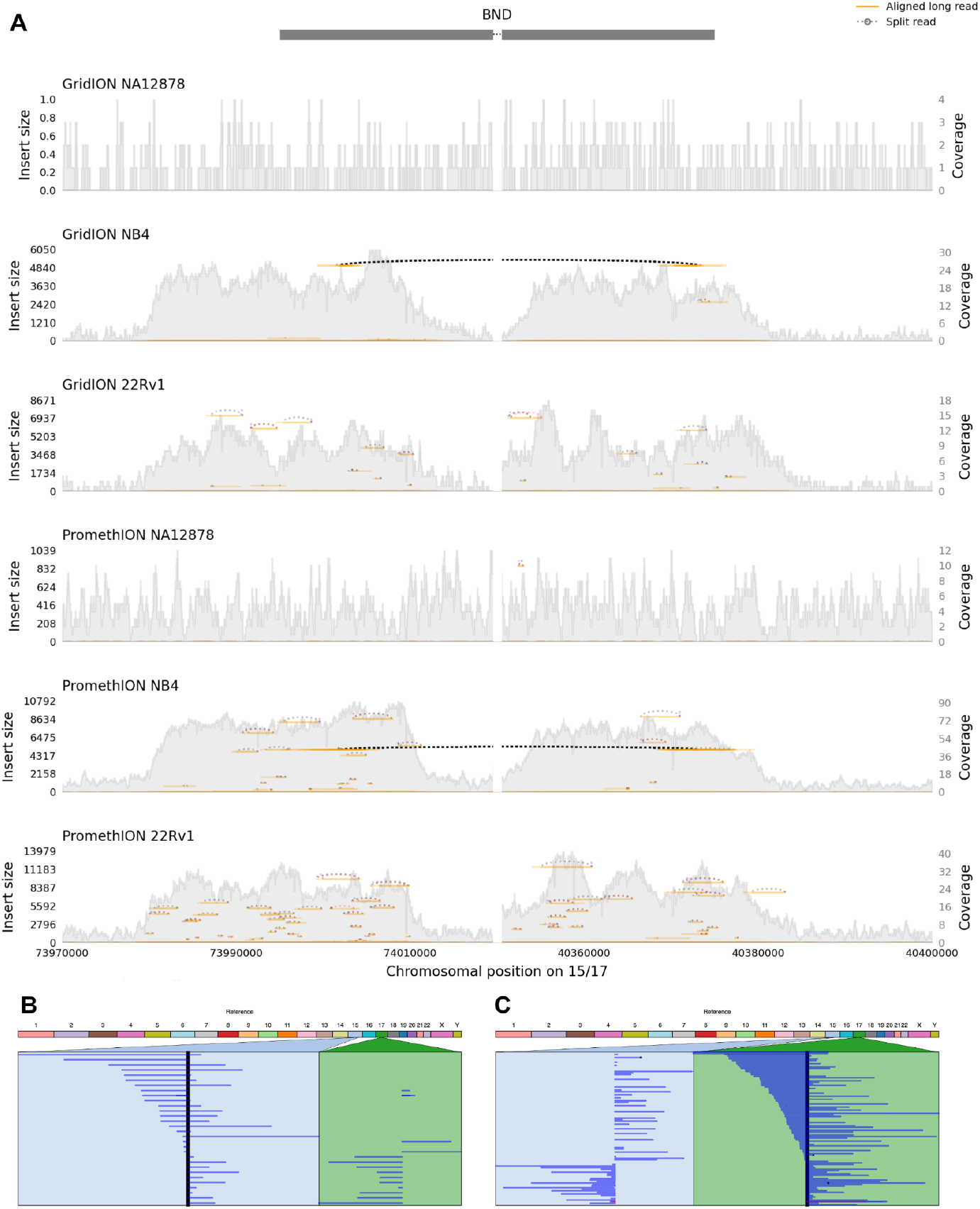
Visualising Structural Variation. A) Using samplot^23^, we visualise reads linking PML (chromosome 15) and the known fusion with RARA (chromosome 17). Only the NB4 sample carries this fusion (indicated by the dashed lines). B), C) Using Ribbon we can visualise individual reads from B) the GridION NB4 sample and C) the PromethION NB4 sample. SVs were identified using CuteSV (supplementary file 1).

When switching to PromethION, the on target read N50 remains similar whilst the rejected read N50 increases to 970 bases. This equates to approximately 2.1 seconds of sequencing time which will reduce the overall efficiency of adaptive sampling. This difference is not explained solely by the change in chunk size which will only add 0.2 seconds to the total rejected read length. Rather this is likely caused by the considerable computational challenge of controlling and addressing 3000 independently sequencing channels in real-time. Notwithstanding, the mean target coverage per sample ranges from 27x to 49.5x. This variation in coverage is, in part, explained by the difference in total target numbers for each sample as well as differences in the input read lengths, library pooling prior to sequencing and copy number variation within the sample itself. To achieve this coverage across three samples without adaptive sampling would require 300-400 Gb of read data or 4 to 5 times the amount we sequenced here. Coverage compared with GridION is significantly improved (Figure 3, D-F) and structural variant analysis clearly identifies the known changes in these samples (Figure 4). These data are also likely sufficient for comprehensive variant analysis.

Finally, we turned to a natural application for adaptive sampling which considers the mappings of rejected reads. Various approaches have been developed using binning of short reads to detect copy number variation by applying a variety of statistical approaches ^14^. These methods also work with nanopore sequencing ^15^, but the resolution of detection will be dependent on the total number of reads generated during a sequencing run. Adaptive sampling increases read count as a consequence of rejecting molecules once they are confidently mapped to an off-target region. We therefore developed a simple approach to bin read counts across the genome such that, on average, each bin would contain 100 reads, and monitored this in real-time using our minoTour tool ^16^. For each barcoded sample changes in copy number are immediately apparent and can be visualised using any change point detection approach, here we use Ruptures (Figure 5) ^17^. As expected, GM12878 (barcode 1) does not show significant copy number changes, whereas NB4 (barcode 2) and 22Rv1 (barcode 3) both closely recapitulate results generated by Bionano optical mapping (Figure 6). As expected, PromethION brings higher resolution to assessment of copy number changes due to the higher total read count (Figures 5,6).

**Figure 5.**
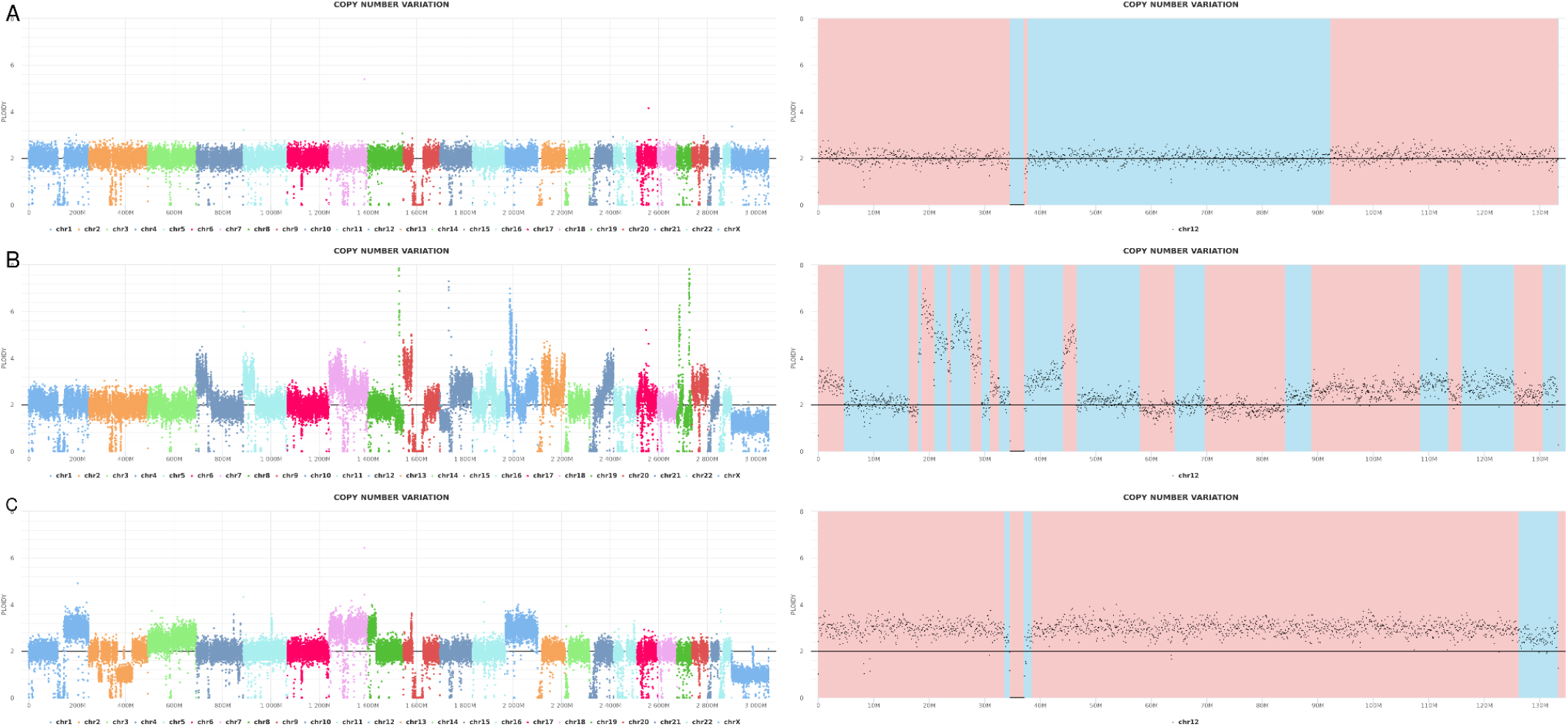
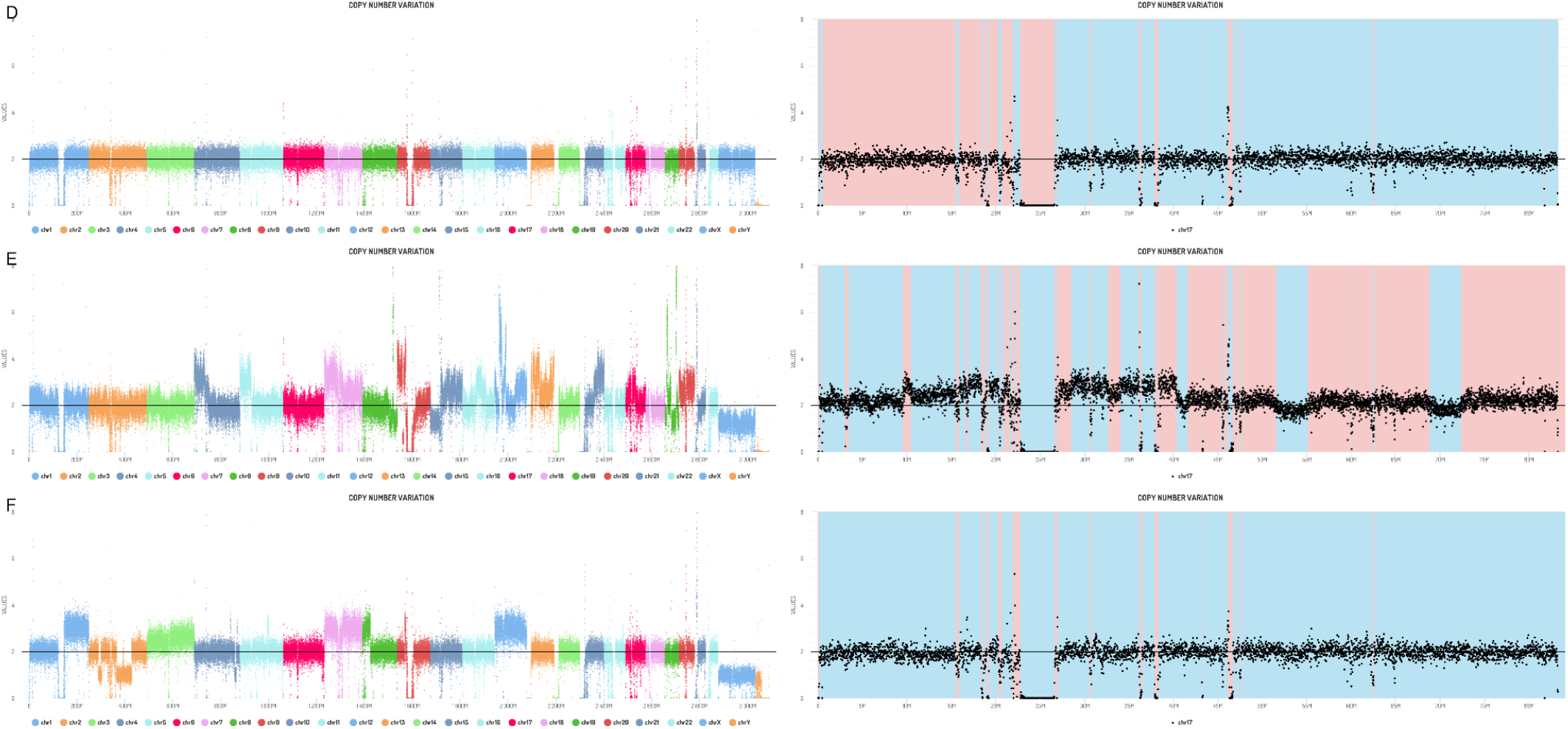
Real-time monitoring of copy number change. MinoTour generates real-time counts of reads dynamically binned such that each bin contains on average 100 reads. Samples A-C shown here mapped to Chm13 T2T reference, with sequence data generated by 72 hours running on a gridION mk1. Samples D-F are mapped to the HG38 reference, with alt chromosomes removed, with sequence data generated by 72 hours running on a promethION. Left hand plots show coverage over all chromosomes, right hand plots show just chromosome 12 for A-C and chromosome 17 for D-F. Red Blue banding indicates change points as dynamically detected by Ruptures. A) barcode 01, GM12878, bin width 86,600 bases. B) barcode 02, NB4, bin width 60,570 bases C) barcode 03, 22Rv1, bin width 76,470 bases. D) barcode 05, GM12878, bin width 19,070 bases. E) barcode 06, NB4, bin width 12,060 bases F) barcode 07, 22Rv1, bin width 20,680 bases.

**Figure 6.**
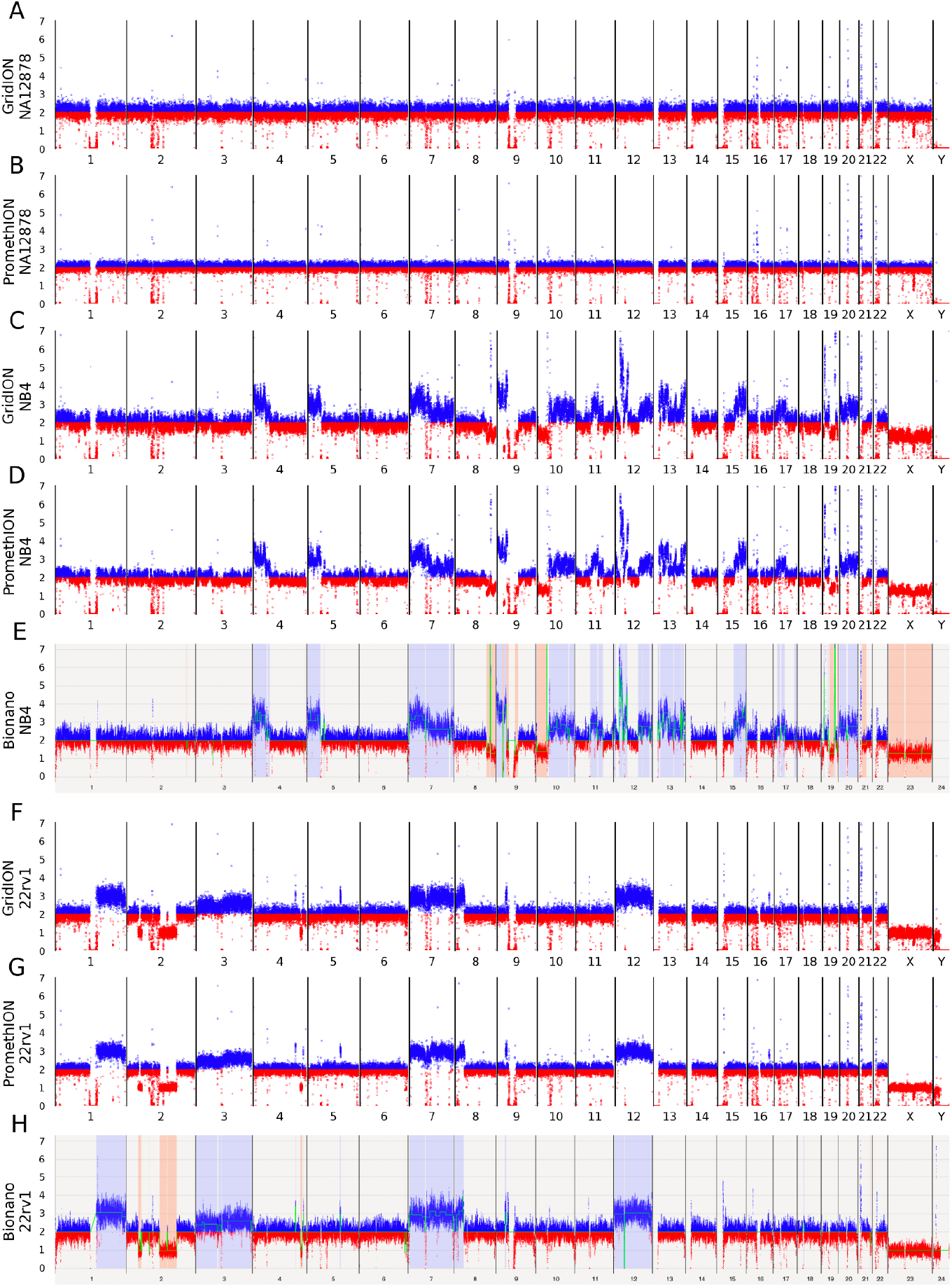
Matched Nanopore Bionano CNV visualisation. Nanopore sequence data from GridION and PromethION compared with Bionano optical reads, all mapped against hg38. Blue points show where binned data indicates greater than expected copy number, red points where binned data indicates lower than expected copy number. A, B) NA12878 showing Nanopore adaptive sampling data only from NA12878 on GridION and PromethION. B, C) NB4 on GridION and PromethION compared with E) showing Bionano optical mapping data for this cell line. F, G) 22rv1 Nanopore adaptive sampling data on GridION and PromethION compared with H) the same cell line visualised using Bionano optical mapping.

## Discussion

Extending readfish to become “barcode aware” enables more sophisticated selection experiments that are better able to exploit adaptive sampling in a variety of contexts. Here we demonstrate that individual samples can be targeted with unique panels of genes, selected based on knowledge of the sample, enabling the user to ask and answer specific questions. On a single MinION flow cell, 3 human genomes can be analysed in real-time with coverage sufficient to detect structural and copy number variation. In this case, yield limitations prevent a realistic assessment of SNPs. However, we anticipate higher yield or running just two human samples per flow cell would enable this. Of course, smaller genomes will generate proportionally higher coverage enabling more samples to be run on a single flow cell as well as providing greater depth for variant calling. However, it is possible to target 3 human genomes on a single flow cell on PromethION devices, to sufficient depth for further SNP analysis. We anticipate further optimisation of the underlying software stack, both within proprietary MinKNOW control software and firmware as well as within tools such as readfish will further enhance yield and throughput, enabling effective targeted sequencing on 6-12 samples on a single PromethION flow cell.

Alongside these targeted experiments, this approach also allows users to simply switch off barcodes for which sufficient data have been generated. This will enable dynamic adjustment of yields obtained from individual samples in barcoded libraries. Our initial testing shows these approaches will work with the full 96 barcodes currently available on nanopore platforms. Coupling multiple samples with barcode aware readfish and real-time analysis of the data obtained will enable faster experimental turn around times, more efficient use of flow cell resources and more comprehensive analysis pipelines.

## Methods

### Synthetic read generation and analysis

To demonstrate our choice of parameters for read mapping and barcode calling, we obtained reads mapping to either chromosome 15 or 17 from the sequenced subset of reads ending up in the pass folder from NB4 (barcode02). Using the ONT Fast5 API (https://github.com/nanoporetech/ont_fast5_api), we generated varying sizes of chunks of signal from the start of these reads incrementing in 0.1 second equivalents (400 samples) to 1 second, then 0.25 seconds to a total of 10,000 samples per read (2.5 seconds). These reads were base called using Guppy (v5.0.16+b9fcd7b) and mapped using minimap2 to the target regions of chromosomes 15 and 17 defined by the trusight RNA fusion panel ^11^. Mapping used minimap2 ^18^ with the -x map-ont and --paf-no-hit option to retain all reads regardless of mapping. We chose the high accuracy model as our truth set as this is the current standard base caller. By using the subset of reads from chromosome 15 and 17 targets only, and hence a smaller reference, we could also test signal based methods for mapping reads including uncalled (v2.2) and sigmap (v0.1) ^3,6^.

For determining alignment accuracy we considered read starts mapping within 50 bases of the truth set as true positives, although for many applications this may be overly stringent. At this stringency, the fast base calling model recovered true mappings with an F1 score of 0.903524 (precision = 0.927901, recall = 0.880395). The code is available in the accompanying data notebooks. As a result we selected 0.8 seconds of data for analysis. Neither sigmap nor uncalled were optimised beyond the default settings and performance could likely be improved further.

For barcoding of data, we used Guppy demultiplexing and tested no other approach. Truth sets were defined using the full length reads as above. We compared the impact of the base caller model on barcode detection and found the fast model recovers the correct barcode with an F1 > 0.9 at 1,600 samples.

### Running readfish Barcoding

Running read until and adaptive sampling requires the ONT Read Until API (version 3.0.0, https://github.com/nanoporetech/read_until_api/tree/release-3.0) and the ONT PyGuppy Client library (version 5.0.13, https://pypi.org/project/ont-pyguppy-client-lib/5.0.13/). Readfish (https://github.com/LooseLab/readfish; commit 9e8794a) was run using a GridION MK1 (MinKNOW v4.3.2; Guppy v5.0.13; minimap2 v2.22), the MinKNOW configuration scripts were configured to serve data in 0.8 second chunks. For PromethION, we ran using a modified version of readfish (https://github.com/LooseLab/readfish; commit c9f5169) on an early release of MinKNOW core 5.1 using a PromethION 24 device (MinKNOW v5.1; Guppy v6.0.6) with the ONT PyGuppy Client library (version 6.0.6, https://pypi.org/project/ont-pyguppy-client-lib/6.0.6/). MinKNOW configuration scripts were left serving data in 1 second chunks for PromethION.

The readfish script carrying out the selective sequencing was “readfish barcode-targets”. This script runs the core Read Until process as specified in the experiment’s TOML file. With a single reference genome the script can select specific target regions on each barcode by using Guppy to base call and demultiplex the raw signal in real-time. The resultant read is then aligned to the reference using minimap2 and is determined to be on or off target depending on it’s barcode assignment and mapping start. For PromethION, we used the mapping returned by minimap2 from within Guppy.

### Library Preparation, Sequencing and Analysis

Barcoded LSK-110 (ONT) sequencing libraries were prepared from either GM12878 cells (Coriell), NB4 cells (gift from M. Hubank) or 22Rv1 cells (ATCC) as described in Jain et al. ^7^. For test experiments bacterial DNA was extracted using genomic tip (QIAGEN). Extracted DNA was sheared to approximately 12 kb using g-Tube (Covaris). All sequencing used FLO-MIN106 R9.4.1 flow cells. Flow cells were run with flushing and reloading as previously described in Payne et al. ^2^.

To investigate structural variation across the dataset, we ran CuteSV on each barcoded sample using standard options but varying the -s MIN SUPPORT values. No SVs in known fusion genes were reported in NA12878 or 22Rv1 (-s 2), known fusions including PML RARA were readily detected in NB4 (-s 5)^13^. SVs were visualised using Ribbon ^19^. To visualise changes in copy number, reads were mapped to hg38, filtered to mapping scores >20 and uniquely mapping. Then the first primary mapping for any read was determined and mappings binned into windows along the genome such that on average each bin contains 100 reads. Runs were monitored in real-time using minoTour (https://github.com/LooseLab/minotourapp/; commit: 1f9c678), providing coverage statistics, mappings and estimates of copy number variation in real-time ^16^. During real-time analysis reads were mapped to Chm13 telomere-to-telomere assembly ^20,21^. Post-run copy number plots were generated using matplotlib with data mapped to hg38 to compare with the output of the Bionano copy number pipeline (see notebooks).

To visualise coverage over specific targets reads were divided into those actively sequenced and those unblocked using the unblocked read ids file generated by readfish. Reads were mapped to hg38, coverage depth calculated using mosdepth v0.3.1^22^ and visualised using matplotlib (v3.4.3).

### Bionano Methods

#### DNA extraction and labelling for Bionano

DNA was prepared from frozen cell pellets of 1.5 million cells using the Bionano Prep SP Blood and Cell Culture DNA Isolation Kit (Bionano Genomics; 80042) according to the manufacturer’s instructions. DNA was homogenised and quantified using Qubit dsDNA BR Kit (Thermo Fisher; Q32853) on a Qubit 4 Fluorometer (Thermo Fisher; Q33238). 750 ng of gDNA was then labelled with Direct Label Enzyme 1 (DLE-1) and DNA backbone stain using the Bionano Prep Direct Label and Stain (DLS) kit (Bionano Genomics; 80005) according to the manufacturer’s instructions. Labelled DNA was quantified using the Qubit dsDNA HS Kit (Thermo Fisher; Q32851) on a Qubit 4 Fluorometer. Labelled DNA was loaded onto a Bionano Saphyr G2.3 chip (Bionano Genomics; 20366) and run on a Gen 2 Bionano Saphyr System (Bionano Genomics; 60325) until 1.320 Tbp of data had been collected for each of NB4 and 22Rv1. This data had respective mapping rates to hg38 reference sequence of 89% and 79%, equating to 382x and 337x coverage respectively.

#### Data analysis

Post run data filtering and analysis was carried out using Bionano Access 1.5.2. For each sample the data set was filtered and sub-sampled to produce 320 Gbp of data with 150 kb minimum length and at least 9 labels per molecule. Filtered data was processed to produce annotated *de novo* assemblies using the default parameters, but with masking using the hg38 DLE-1 SV Mask BED file. Structural variant (SV) and copy number variants (CNV) coordinates were then visualised using Bionano Access. All described analysis was performed on dedicated Bionano compute with the following versions installed: Bionano Access1.5.2, Bionano Tools 1.5.3, Bionano Solve Solve3.5.1_01142020, RefAligner 10330.10436rel, HybridScaffold 12162019, SVMerge 12162019, VariantAnnotation 12162019, Compute on Demand 1.5.1.

## Acknowledgements

The authors thank Mike Hubank and Nigel Mongan for gifts of cells and useful discussions. This work was supported by BBSRC iCASE studentship awards to RM and AP. In addition we acknowledge funding from the BBSRC (BB/N017099/1) and Wellcome Trust (grant number 204843/Z/16/Z).

